# Implementing QR Codes in Academia to Improve Sample Tracking, Data Accessibility, and Traceability in Multicampus Interdisciplinary Collaborations

**DOI:** 10.1101/2021.03.22.436472

**Authors:** Cristian Hernandez, Elham Aslankoohi, Pavel Frolikov, Sri Kurniawan, Marco Rolandi

## Abstract

With the growing number of multicampus interdisciplinary projects in academic institutions, there is an increasing need for tracking systems that make device and sample data and associated results instantly accessible to all collaborators involved. This need has become particularly salient with the COVID pandemic and consequent travel restrictions that have hampered in person meetings and laboratory visits. We report a Quick Response (QR) code tracking system that integrates project management tools for seamless communication and tracking of materials and devices between three manufacturing sites (cleanrooms) and six research laboratories across five universities. We use this system to track the design, fabrication, and quality control of bioelectronic devices (in the cleanroom and engineering laboratories), *in vitro* experimental results (in a biology laboratory), and *in vivo* testing (in the school of medicine). Setting up a tracking system for multisite interdisciplinary teams improves traceability, efficiency, and the quality of produced results making it easier to accomplish project milestones on a tight timeline. This tracking system is particularly useful to track device issues and ensure engineering device consistency when working with expensive biological samples *in vitro* and animals *in vivo* to reduce waste of biological and animal resources associated with device failure.

## Introduction

Modern research encourages multicampus collaboration amongst different disciplines to share concepts and methods as well as to answer more complex questions that cannot be answered within a single discipline [1]. If multicampus interdisciplinary research is not executed properly the produced results and milestone completion success rate can be jeopardized due to inadequate project coordination. One of the biggest challenges of project coordination is communication between experts across different fields that often do not share terminology and tools. This communication is critical when devices and samples are transferred between different laboratories. For example, it is necessary for researchers in one laboratory that need a device for their experiments to receive the proper protocols and understanding of the device’s functionality from researchers that fabricated and tested the device in a different laboratory. This exchange happens often in bioengineering disciplines in which materials and devices are typically developed in an engineering laboratory and are used in a biological or medical setting [2] such as in tissue engineering, [3, 4] medical devices and sensors,[5] and bioelectronics[6-8]. Typically, this exchange of information happens in person via exchange of personnel between laboratories so that collaborators can understand and trust each other through socialization [9]. However, the recent COVID pandemic has not only hampered travel and visits across different laboratories and campuses, but has forced most researchers in the same laboratory to work alone due to limited room capacity restrictions. This limited interaction has made successful materials and sample exchange more and more challenging with high risk of failed experiments due to incomplete communication and lack of device traceability between collaborators.

To mitigate some of these challenges, project management with a tracking system for devices and data can allow interdisciplinary teams to perform quality work while reaching task milestones on time and staying within budget [10, 11]. For example, QR codes have become a common tool to share information and are easily accessible with any modern smartphone [12]. In academic settings, QR codes provide data on classroom links, tutorials, vCard contacts, uniform resource identifiers, e-mail addresses, map directions, text, and chemicals. And have started to surface in the research environment for quick tracking of biological specimens [11][13]. However, current use of QR codes in academic research remains limited.

Here, we introduce a QR code tracking system for materials and devices as a solution to improve project coordination across five collaborating universities (UC Santa Cruz, University of Montana, University of Utah, Tufts University, and UC Davis) during the COVID pandemic as part of a DARPA multicampus project associated with the BETR program. This project involves the use of bioelectronic sensors and devices [14, 15] combined with AI algorithms to speed up wound healing[16]. Bioelectronic devices are designed at UC Santa Cruz; fabricated with up to ten clean room processes at University of Montana, University of Utah, and UC Santa Cruz; tested at UC Santa Cruz; and then shipped either to Tufts University for *in vitro* experiments or to UC Davis for work *in vivo*. Inspired by Toyota’s Total Production System[17], we introduced device QR codes as a device “Kanban” to solve two main issues: (1) inventory management to make sure devices are ready “just in time” when the biological experiments are scheduled so that we avoid poor performance due to device age or missed experiments due to delays in fabrication, and (2) the need to trace the root cause of device failure during biological experiments to specific steps in device design or fabrication.

## Materials and Methods

### QR Code Generator

We use the outsourced management tool desktop app, https://www.qr-code-generator.com/, that generates QR codes. The project management tool provides dynamic functionalities allowing the codes to be edited even after having been printed onto a label. This capability is useful for devices that go through various fabrication steps and quality control checkpoints before completion. It allows the fabricator to make updates throughout the products lifecycle without having to continuously reprint the QR Label. The QR code generator allows the user to customize the information: QR code name, basic information, categories, and contact information displayed on the scanning phone. We use a unique identifier known as a batch number to name the QR code. Batch numbers keep track of devices that are built in bulk. This study requires assigning batch numbers to various wafers fabricated at the same time. A wafer yields seven devices once completed. The batch number consists of nomenclature that provides relevant information on the wafer’s fabrication. The batch number’s nomenclature, 123456-7-8-ABC-9, represents the following:

1. 123456: The digits provide the date that manufacturing started on the wafer.
2. 7-8: The digits identify the number of wafers that started fabrication on the same date.
3. ABC: The letters are abbreviations that provide the device’s design name.
4. 9: The final digit conveys the device’s design version.

At a certain step of the fabrication process, wafers are diced into individual devices and each device is assigned a part number. The part number’s nomenclature is the batch number but with an added letter at the end ranging from A-G to represent the seven devices, for example123456-7-8-ABC-9-D. Without these two unique identifiers the devices would not be traceable with the QR code tracking system. The user interface provides the following customized categories when scanning the label: (1) batch information, (2) wafer build process, (3) device build process, (4) batch tracking, (5) quality control, (6) shipping schedule, and (7) sample testing. A label maker prints the QR code and batch number. A one-inch label goes on a plastic container that protects the wafer throughout its lifecycle. The wafer goes through fabrication, quality control and is diced into individual devices. The fabricators assign each device with a new unique identifier that is known as a part number. The individual devices continue fabrication and quality control until completion and get shipped to the collaborators for *in vitro* testing. During a device’s production lifecycle, all the categories are updated manually in the QR code generating desktop app and Asana.

### Desktop and Mobile

We developed at UC Santa Cruz an application that integrates data from QR codes onto Asana. The application mirrors functions from the outsourced QR code generator, but allows the user to make updates on a mobile phone and automatically transfers the data onto Asana. The application is opensource and will provide users an opportunity to integrate the application into their projects for free or to use the outsourced QR code generator that this study first started using to track device data. The application is further explained in the results section.

## Results

We used outsourced project management tools, QR code generator and Asana, to track all bioelectronic device production lifecycles to ensure engineering device consistency to reduce waste associated with device failure (Fig 1). The QR generator generates a QR code and assigns a batch number for any wafer and tracks the wafer throughout its lifecycle. The researcher can scan the QR label at any time during the products lifecycle to access any information needed to make a decision (Fig 1a).The QR code consists of different parts that correspond to different steps of the process such as batch info, wafer build process, device build process, batch tracking, quality control, shipping schedule, and sample testing (Fig 1b). The

**Figure 1.**
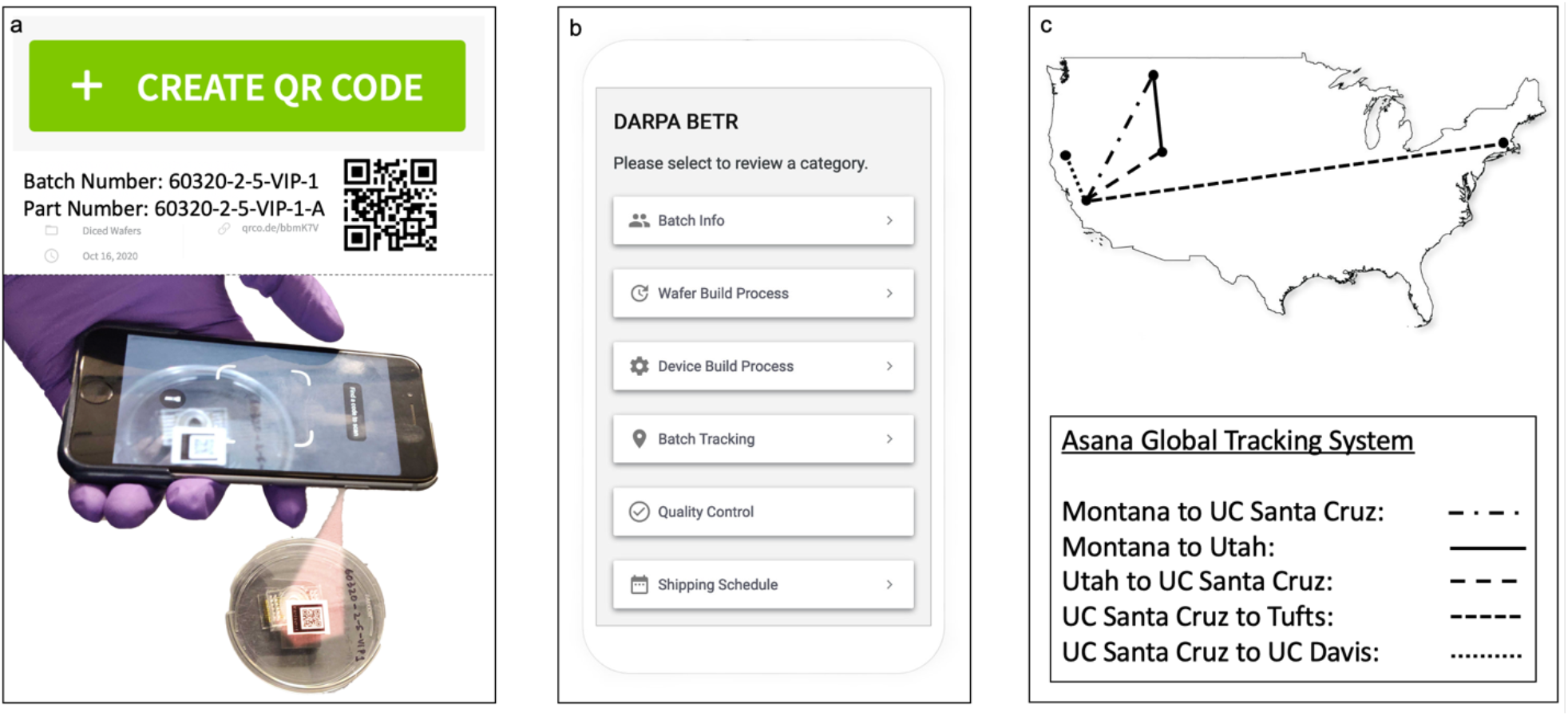
Tracking System Overview: (a) Generate QR Code on subscription-based desktop application and label device container with unique identifier and QR Code. Scan QR Code to access data. (b) Data accessed focuses on the following categories: batch info, build process, batch tracking, quality control, shipping schedule, and sample testing. (c) Data from the QR codes tracking system is implemented into Asana, a global tracking system that we customized to provide device data traceability amongst five universities. Asana tracks the location and transit of the device across the different collaborating institutions. This tracking is important for inventory management to make sure that there are always enough devices at each step and location so that the subsequent fabrication step or experiment is not delayed.

In the first step of the tracking process, the researcher that fabricates a device also prints a QR code with a unique identifier (fig 2.A). The batch number identifies and traces the bioelectronic device production history and relevant issues that occurred during manufacturing and testing. The nomenclature provides instant information on the wafers to determine the following: date of manufacture, wafer identifier, number of wafers in production, design identifier, and design version. For example, 92220-4-4-VIP-1 allows our fabricators to determine wafer fabrication started on September 22, 2020, it is the fourth wafer out of four batches produced, and it is the first design version of the Vertical Ion Pump, which is a bioelectronic device design. The devices are tracked by placing a label on the plastic containers that secure and protect the devices throughout production (fig 2.B). The outsourced project management tool allows the researchers to generate and update a QR code (fig 2.C). QR codes are generated by selecting the green button titled “Create QR Code.” This user-interface then provides a customizable layout that can be edited and reedited with data pertaining to the needs of the team.

**Figure 2.**
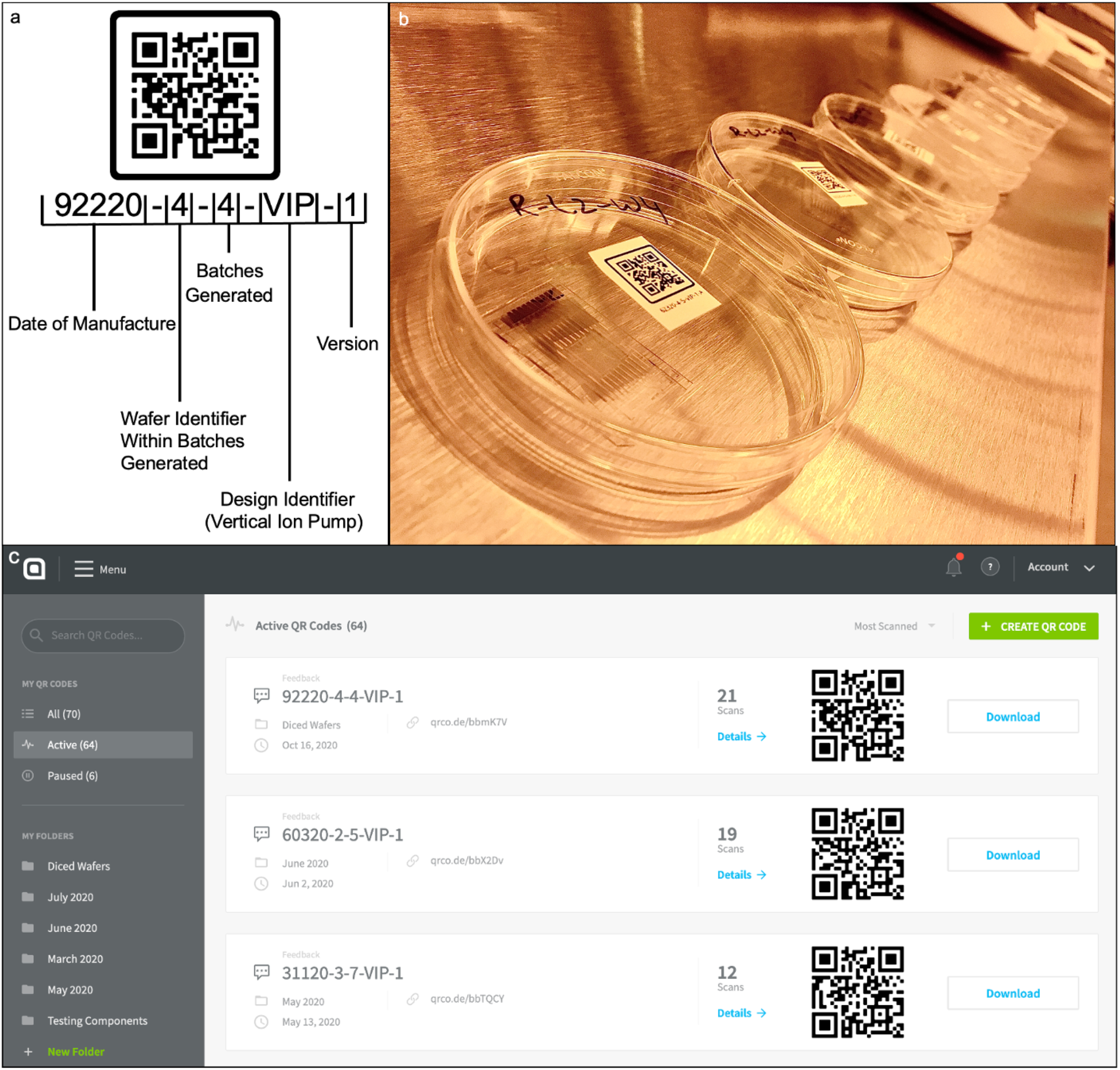
QR Code Tracking System: (a) A detailed breakdown of the unique identifier’s nomenclature that is tied to a QR Code, and (b) label device container with unique identifier and QR Code. (c) User interface subscription-based desktop application used to generate QR Code.

The QR code system tracks data that has been separated into categories to keep the information organized (fig.3). Batch info provides the researcher with general data such as batch number, initial fabrication location, substrate material, fabricator name, etc. (fig 3.a). For example, fig 3.a lets the researcher know that wafer 92220-4-4-VIP-1 was initially fabricated in the Montana cleanroom on September 22,2020. The wafer substrate is Borofloat 500 um. The device has been shipped to Utah to continue more fabrication and was diced into seven devices that received part numbers. The wafer build process consists of six steps and the system displays the wafer’s progress through the fabrication process (fig 3.b). Completing the first six steps leads to the wafer being diced into seven devices that receive new part numbers. The system tracks the fabrication process of each diced device and if any failures occur during a specific step (fig 3.c). The wafer and the devices it yields after dicing are tracked throughout fabrication and testing to determine which collaborators have come in contact with the device(s) (fig 3.d). The system tracks quality control performed on the devices to determine the cause of failure and at which step during production the device(s) failed (fig 3.e). Shipping for the device(s) is tracked to help researchers plan for fabrication or testing based on the package’s arrival (fig 3.f). As the project continues, more categories will be added. This tracking is important so that if any issues occur during *in vitro* testing, the researchers are able to track down all of the specific processes and dates that that wafer has undergone. For example, if there were any issues with bonding two of the wafers together that would result in leakage of fluid from the device, the QR code would allow the UC Santa Cruz team to trace the wafer back to a specific device process and batch in the cleanroom in Montana. If leakage were to occur in most of the devices produced with the same process on a specific day, then we will determine that we required to optimize the process so that the bonding between the wafers is more robust.

**Figure 3.**
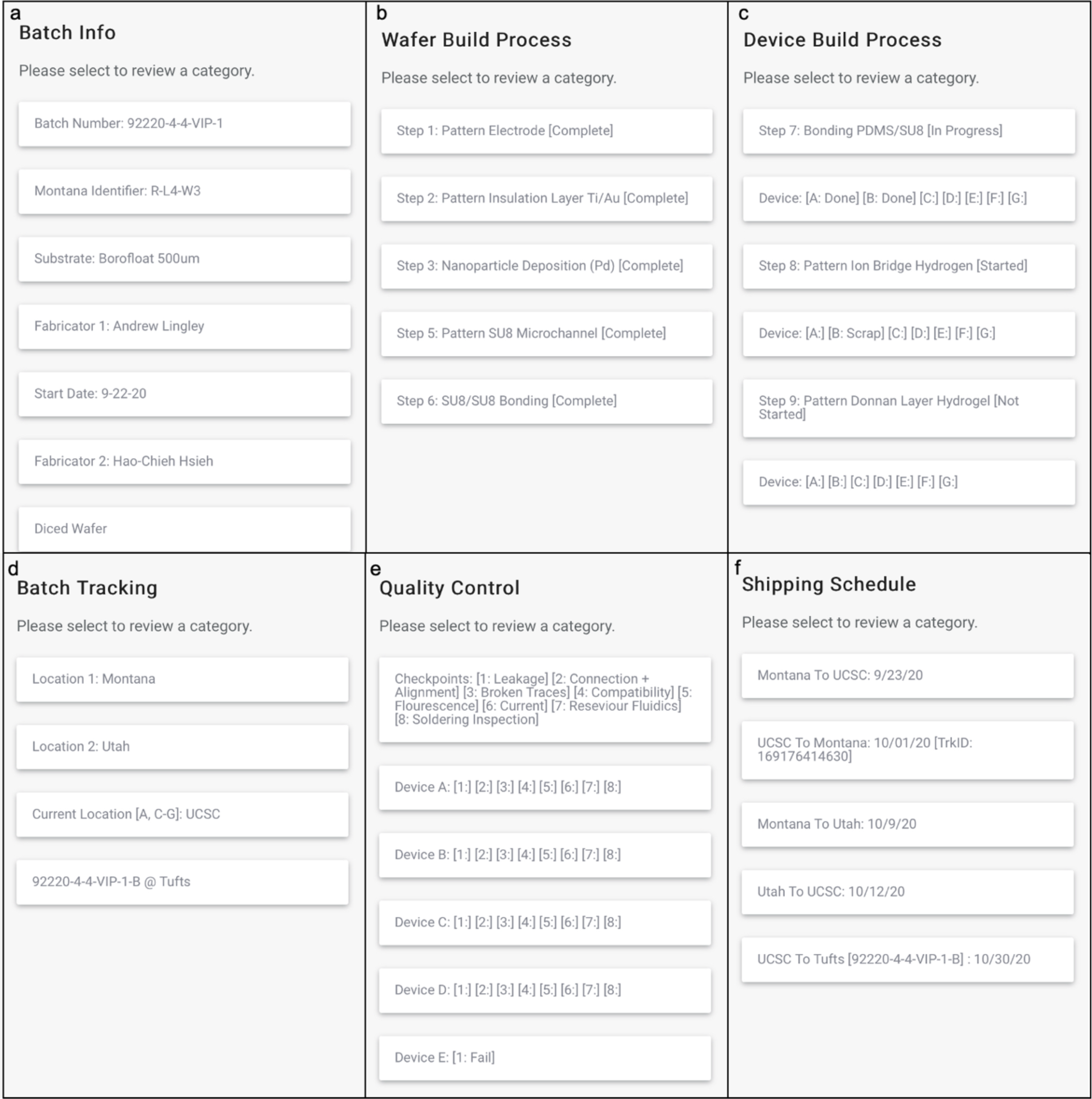
Categoric Data Points that QR Code System Tracks: (a) Batch Information, (b) Wafer Build Process, (c) Device Build Process, (d) Batch Tracking, (e) Quality Control, & (f) Shipping Schedule.

Using the QR code generating tool, a user can scan the QR code and access the data via mobile phone, the data can only be updated on a desktop website application. All the information then has to be manually inputted into Asana for all the collaborators to easily access and trace data in one organized location. To provide more real time updates, prevent any loss of data, simplify data implementation, and automatically transfer data from the QR code tracking system to Asana, we developed a mobile application (Fig. 4). This application is open-source and will be made available to all interested universities to help academic researchers manage their data better. Fig. 4.A displays the user interface of the application. With this application, a user can make updates on the mobile and desktop application. The application uses py4web (a database driven web framework) for the backend framework. The QR Code tracker website is a user navigable database so py4web is a perfect fit as the framework. The user interface for the tracker website was specifically designed to mimic the outsourced QR code generator system with added accessibility. A user can create their own categories and fill them with any data that mitigates project issues directly on a mobile phone. The tracker website uses very little computational power because it uses pythonanywhere.com for hosting. The biggest challenge with the application development came from integrating the Asana API. The application provides a functionality to automatically transfer all the data on the QR code tracking system into Asana by selecting the “Update Asana” button.

**Figure 4.**
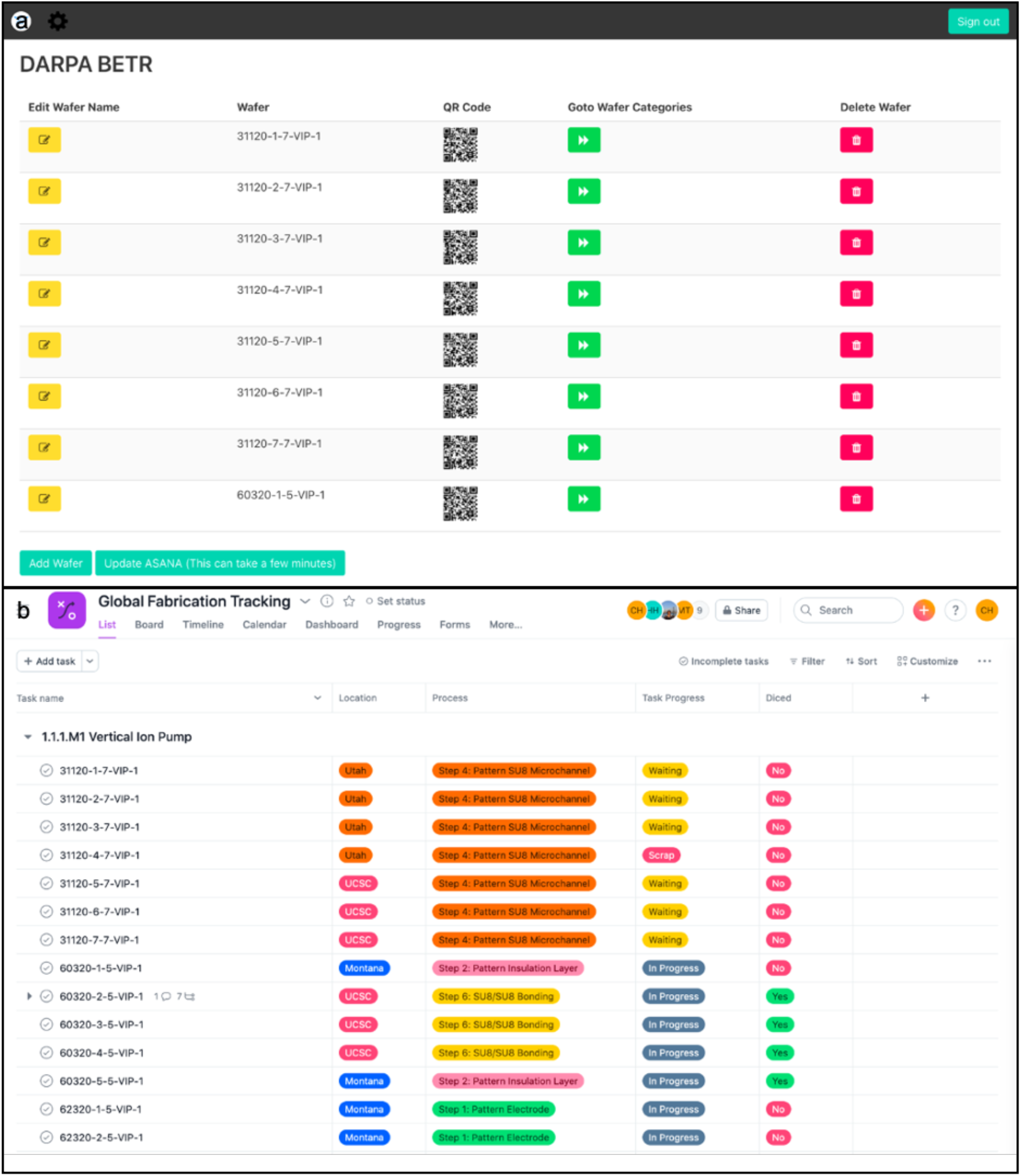
(a)Application Tracking System that Integrates QR Codes and Asana, (b)Asana Global Tracking System

Fig 4.B displays a snippet of the data that is automatically transferred into the project management tool Asana when using the application. All the data that is stored and shared between the researchers who focused on fabrication and testing gets added onto Asana. This allows the data to be easily understood and provides easier access to all users across three universities. Ten fields are filled with data points that allow the users to coordinate the project accordingly. The fields are location, assignee, due date, process, task progress, diced, expected devices, devices yielded, scrap percentage, and testing. Location conveys which university the device(s) are at. Assignee provides the name of the researcher that is in charge of the device(s). Due date is the date the device(s) must finish in order to reach milestones in a timely manner. Process provides the fabrication step the device(s) are currently going through. Task progress details if the device(s) are in progress, not started, done, waiting, or scrap. Diced details if the wafer has been cut into seven individual devices. Expected devices provides the number of devices the wafer needs to generate when finished with fabrication. Devices yielded provides the number of devices that were generated after fabrication completion. Scrap percentage is a basic formula that provides the percentage of failure from a wafer. The formula is ([scrap device(s) / expected devices] * 100). The goal is to keep device failure below 25%. Knowing the scrap percentage opens up the dialogue on what caused the failure and how can it be prevented. This feature is provided to understand the causes of failure in the first prototype and the hope is that the knowledge will mitigate any risks from the same failure occurring in future fabrications. Testing details if the device will undergo in vitro or in vivo testing.

In order to track inventory at different steps, we curate the data from Asana into a table that is shared with the principal investigators of the project (Table 2). This table allows all collaborators to know that the Montana cleanroom has delivered 63 devices out of 140 that were ordered. The Montana team plans to start fabrication on 28 devices. The Utah cleanroom has 49 devices in production with 35 on step 1 and 14 on step 2. UCSC cleanroom has multiple devices in production at various steps. This table allows for the researchers that perform device fabrication to organize their fabrication runs to supply the demand for devices for *in vitro* and *in vivo* testing. In turn, it allows for the researchers performing *in vitro* and *in vivo* testing to plan and prepare their experiments accordingly to when the devices will be available.

**Table 2:** Total quantity of devices in production and current fabrication steps

| Process |  | Montana | Utah | Santa Cruz |
| --- | --- | --- | --- | --- |
| Devices Ordered |  | 140 | N/A | N/A |
| Devices Delivered |  | 63 | 0 | N/A |
| Devices Currently In Production |  | 28 | 49 | 136 |
| Step 1 | 3 days | 0 | 35 | 0 |
| Step 2 | 1 day | 0 | 14 | 77 |
| Step 3 | 4 days | X | X | 28 |
| Step 4 | 1 day | 0 | 0 | 28 |
| Step 5 | 1 day | 0 | 0 | 28 |
| Step 6 | 4 days | 0 | 0 | 28 |
| Step 7 | 1 days | X | X | 3 |
| Step 8 | 2 days | X | X | 5 |
| Step 9 | 2 days | X | X | 0 |

## Conclusion

Multicampus interdisciplinary projects need a QR code tracking system and can stem to benefit in project coordination especially during Covid-19. Here, we describe a QR code tracking system that we used to track device design, fabrication, testing, and *in vitro* and *in vivo* use across five campuses as part of a DARPA funded multidisciplinary project. This system allows for more efficient “just in time” inventory management and faster identification of root cause of failure by tracking the history of a specific device. We believe that this system can help these larger projects achieving a more efficient use of resources by limiting the number of failures. This is particularly important for bioengineering projects that involve engineering devices in biological and animal use to reduce waste of biological and animal resources. Finally, we hope that improved efficiency may increase the yield of results from federally funded research.

## Acknowledgment

This research is sponsored by the Defense Advanced Research Projects Agency (DARPA) through Cooperative Agreement D20AC00003 awarded by the U.S. Department of the Interior (DOI), Interior Business Center. The content of the information does not necessarily reflect the position or the policy of the Government, and no official endorsement should be inferred. The funders had no role in study design, data collection and analysis, decision to publish, or preparation of the manuscript. The authors thank Harika Dechiraju and John Selberg for valuable feedback and discussions.

